# Biomedical researchers use preprints for speed, but hesitate to credit them in evaluation

**DOI:** 10.64898/2026.03.02.709147

**Authors:** Xi Hong, B. Ian Hutchins, Chaoqun Ni

## Abstract

The preprint ecosystem has expanded rapidly over the past decade, fundamentally altering science communication. Yet, the scholarly community’s attitudes toward this shift remain underexplored. Through a large-scale survey of US and Canadian biomedical scholars, we provide a comprehensive analysis of preprint utilization, perceived impact, and integration into academic credit systems. We find robust engagement across reading, citing, and submitting preprints; however, this activity is driven primarily by a desire for rapid dissemination rather than a foundational commitment to open science. Furthermore, while preprints are valued as networking assets, perceived career penalties during formal academic evaluations stifle broader cultural adoption. Crucially, to navigate the absence of formal peer review, scholars report a heavy reliance on author reputation as a primary heuristic for evaluating a preprint’s credibility and guiding their reading and citation decisions. Notably, despite acknowledging preprints’ role in accelerating knowledge sharing, scholars express significant concerns regarding fraud and misinformation, particularly amid declining public trust in science and emerging threats to scientific integrity from artificial intelligence. To resolve these tensions, the preprint ecosystem must evolve beyond prioritizing speed to foster genuine academic dialogue. Simultaneously, evaluation frameworks must adapt to the realities of preprinting, and innovative quality-control mechanisms are urgently needed to balance rapid dissemination with rigorous scientific integrity.

## Introduction

Funding agencies, research institutions, and individual scientists all seek mechanisms to accelerate scientific discovery; posting complete findings as preprints, avoiding the one-year delay that journal peer review adds to scientific communication (Björk & Solomon, 2013), has been proposed as a simple way of accelerating science (NIH, 2017). In recent years, there has thus been growing attention to preprints as an increasingly visible component of scholarly outcomes (Abdill et al., 2020). Preprinting is particularly attractive because it does not require substantial additional funding, and may be achievable through the existing incentive structures in the research ecosystem. In biomedical fields, preprints have grown rapidly. Launched in 2013, the primary biomedical preprint server, bioRxiv, has posted over 310 thousand preprints, receiving over ten million views per month (Sever et al., 2026). The role of preprints became important during the 2020 pandemic (Sever et al., 2026). In the first 10 months of the pandemic, preprint servers hosted nearly 25% of pandemic-related studies (Fraser et al., 2021). Preprints substantially altered both the timing of scientific communication and its governance by eliminating the peer-review process that scientists have traditionally relied on for quality control.

Scientists and other decision-makers have different motivations for posting preprints. Common reasons scholars submit preprints include establishing priority for their findings (Desjardins-Proulx et al., 2013), disseminating findings early and quickly (Chiarelli et al., 2019), obtaining recognition for their research efforts, receiving open and public feedback (J. A. Teixeira da Silva, 2021), sharing new knowledge with peers and the public (Schloss, 2017), and accelerating the pace of science (Kaiser, 2017). Empirical studies documented a positive relationship between submitting preprints and higher citations and social impact of the later peer-reviewed published versions (Chen et al., 2017; Fraser et al., 2020; Fu & Hughey, 2019; Metcalfe, 2006; Wang et al., 2020). This indicates that there is measurable impact of sharing results earlier. Recognizing this potential, funders have also begun to formalize the use of preprints in their funding processes. NIH, the Chan Zuckerberg Initiative, and the European Research Council have launched policies to encourage preprints (Puebla et al., 2022). Thus, preprints have a demonstrated appeal attributable to multiple motivations that have led to increased adoption. However, these motivations relate primarily to posting preprints, rather than reading, citing, or evaluating preprints. Trust is critical for the success of preprints in accelerating science.

Tensions between rapid dissemination and concerns about quality, credibility, and academic recognition may thus constrain the ecosystem’s further development. Concerns about the quality of research without peer review have discouraged scholars from posting preprints (Fraser et al., 2022). In South Korea, despite recognizing preprints’ contributions to promoting open access, rapid feedback, and earlier citations, most respondents still had a negative attitude toward preprints due to concerns about scientific integrity (Yi & Huh, 2021). A survey in South Asia showed that among scientists who had read preprints, only 64% trusted them (Das & Gutam, 2020). The question of preprint reliability has led researchers to study the differences between preprints and their published versions (Carneiro et al., 2020; Janda et al., 2022; Nelson et al., 2022; Yin et al., 2026). Some found that there were evident differences in titles, data, or conclusions between preprints and their published versions (Gehanno et al., 2022). Moreover, attitudes toward preprints vary among policymakers. One study reported that among the 383 Asian academic society journals, only 7% accepted manuscripts that had previously been submitted as preprints, and 2% allowed preprints as references (Choi et al., 2021). Another study suggests that decision-makers in academia may still view preprinting culture conservatively, continuing to rely on traditional peer-review-based metrics in tenure and promotion decisions (Chiarelli et al., 2019). These results show that despite their potential, preprints are still viewed with notable distrust by some users and evaluators.

Because the value of the preprint ecosystem depends on how these views shape scientists’ subsequent decisions to read, cite, submit, and evaluate preprints, it is essential to comprehensively understand researchers’ attitudes toward and perceptions of preprints. The literature offers relatively limited large-scale empirical evidence capturing the use and perceptions of preprints across multiple academic roles. Most prior studies focus on specific actions, mainly scholars’ behavior in either reading or submitting preprints, but not both. The literature still lacks integrated evidence to examine different actors’ behavior of using preprints, alongside their perceptions of preprints’ quality, career impact, and credit in academic evaluation, and the complex interplay between these behaviors and perceptions. Besides, existing evidence on this topic is more frequently drawn from Asian countries such as India, South Korea, and China, which suggests the need for further evidence from other regions. Given the central tension that these collective studies identify—a promising avenue to accelerate science versus a risky science communication strategy that may misinform readers—it is imperative to identify scientists’ multifaceted perceptions about preprints as authors, readers, and evaluators. These concerns point to a need for systematic study of the perceived strengths, limitations, and incentive structures surrounding preprints.

Our survey aims to address these gaps by providing a comprehensive view through a large-scale survey of US and Canadian scholars in the field of biomedicine. It contributes to this topic by examining scholars’ overall use of preprints and their underlying motivations, perceptions of preprint quality and impact, and the credit assigned to preprints in academic evaluation.

## Data and methods

### Survey design and distribution

This study uses a survey of scholars in the field of biomedicine in North America to examine the adoption and perceived impact of preprints. The core questions of the survey are organized into three sections: (1) familiarity with and use of preprints, including the frequency of and the reasons for reading, citing, and submitting preprints; (2) perceived quality and impact of preprints, examining respondents’ attitudes toward the overall quality of preprints and their impact on the scientific community, society, and scholars’ professional development; (3) credit given to preprints as research achievements, focusing on how respondents perceive preprints being credited in grant applications, faculty hiring and tenure promotion.

To distribute the survey, we compiled email addresses from academic papers and sent the survey via Qualtrics to the corresponding addresses. Specifically, a dataset of 13,043,450 publication records from *iCite*, the NIH Open Citation Collection (Hutchins & Santangelo, 2025), was matched to the Web of Science database using the Digital Object Identifiers (DOIs) to retrieve the authors’ email addresses. The dataset was further restricted to works published since 2017 and to authors affiliated with institutions in the United States or Canada. This process yielded 281,940 unique email addresses, including 243,627 from 1,079 U.S. institutions and 38,313 from 91 Canadian institutions. The survey was distributed through Qualtrics from March 13 to May 22, 2025. In total, 68,600 emails bounced or failed to deliver. Of the 213,340 emails successfully delivered, 5,855 recipients initiated the survey, corresponding to a 2.7% response rate. Ultimately, 1,874 respondents completed the survey, indicating a completion rate of 32.0%. This study was approved by the affiliated university’s Institutional Review Board, and all participants provided informed consent prior to participation.

**Table 1.**
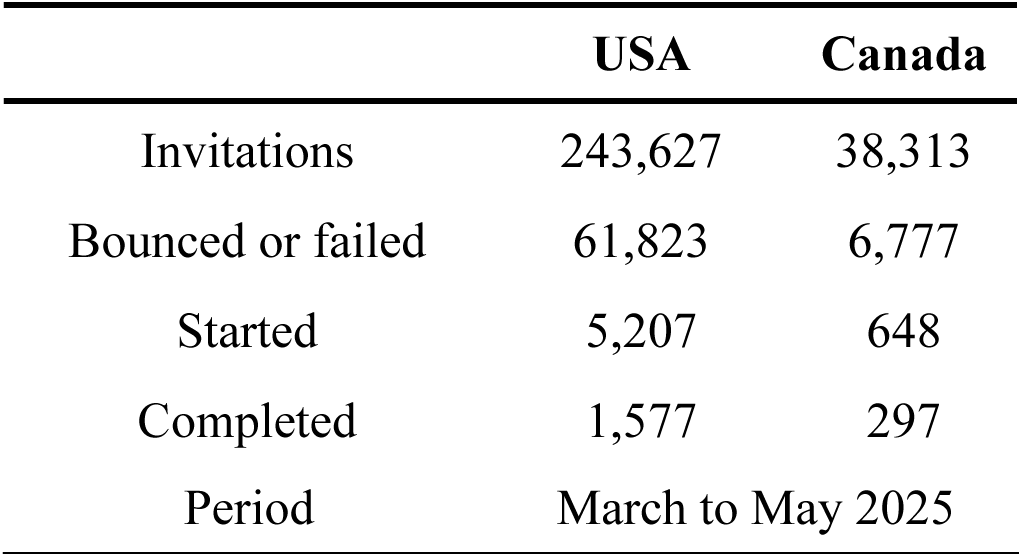
Survey data collection.

### Analytical sample description

After removing invalid responses, such as those with 95% of the questions left unanswered, the final analytical sample for this study includes 1,757 participants. Specifically, researchers from the United States constitute 83.7% of participants, followed by Canada (16.3%). Among the respondents, 76.7% are in the sector of higher education and research institutions, in which 53.3% are full professors, 25.4% are associate professors, and 8.6% are assistant professors. Respondents in other academic ranks include research faculty (5.4%), postdoctoral researchers (2.5%), PhD students (1.2%), teaching faculty (0.6%), and other roles (3.0%). Of the respondents, 61.9% describe their gender as male, while 36.0% identify as female and 2.1% as others. In addition, 80.3% of participants report being White or Caucasian, followed by Asian or Pacific Islander (9.2%), other races (5.4%), Hispanic or Latino (3.3%), and Black or African (1.9%).

### Statistical approaches

We examine scholars’ use and perceptions of preprints through post-stratification-weighted descriptive analysis and ordinal logistic regressions. To mitigate the sample limitation in representing the population, we constructed post-stratification weights based on eight strata defined by the joint distribution of four fields and two countries (see Supplementary **Note 1** for details). Respondents were then weighted to match the corresponding distribution in the population of email invitees. The outcome variables include the frequency of reading, citing, and submitting preprints; respondents’ reasons for (not) engaging in these practices; perceived quality of preprints; perceived impact of preprints on the scientific community, society, and scholars’ professional development; and the perceived credit that preprints should receive as scholars’ representative research achievements in academic evaluation. Moreover, we compare preprint use and perceptions across groups within each country, field, and academic rank through ordinal logistic regressions (see Supplementary **Note 2** for details).

In addition, we apply deductive content analysis (Elo & Kyngäs, 2008; Krippendorff, 2004) with sentiment classification to identify the themes and classify sentiment in respondents’ free-text comments (N = 424) on preprints. Using a structured three-theme framework: perceived quality of preprints, perceived impact of preprints on the scientific community and society, and perceived impact of submitting preprints on scholars’ professional development, we assign each comment zero to three theme labels based on its semantic meaning, along with a sentiment classification (negative, neutral, or positive) based on its sentiment tone.

## Results

### Overall popularity of preprints

The survey results show a generally active engagement with preprints among scholars in multiple academic contexts, including reading, citing, submitting, and using them as representative achievements in grant applications, faculty-job applications, and tenure promotion. As shown in Figure **1A**, 56.3% of respondents reported reading between 1 and 5 preprints per month over the past two years, while 6.5% read between 6 and 10 and 3.5% read 11 or more. The remaining 33.7% reported not reading any preprints. When asked about preprint submissions over the last two years, 43.0% of participants reported submitting between 1 and 5 preprints, while 49.6% had not submitted any. A smaller proportion submitted more extensively, with 5.9% submitting 6 to 10 preprints and 1.5% submitting 11 or more. In addition, 28.8% of respondents reported citing preprints in their research, while 71.2% indicated they had not (Figure **1B**). Among respondents who had grant application experience in the past two years, 25.7% included preprints in their grant applications, while 74.3% did not. Such use in grant applications is echoed by reviewers’ experience: among participants who had served as grant reviewers, 37.0% reported that they had never reviewed proposals that included preprints as part of the principal investigator’s research achievements, while 33.2% stated they had rarely encountered this (Figure **1C**). Another 23.2% indicated they sometimes saw preprints used, whereas only 6.6% reported frequent exposure. Last, among those who had served on tenure, hiring, or promotion committees, 32.6% reported that they had sometimes encountered candidates listing preprints as research achievements, while 27.1% had never seen it, 28.7% reported rarely encountering it, and 11.6% reported frequent experience.

**Figure 1.**
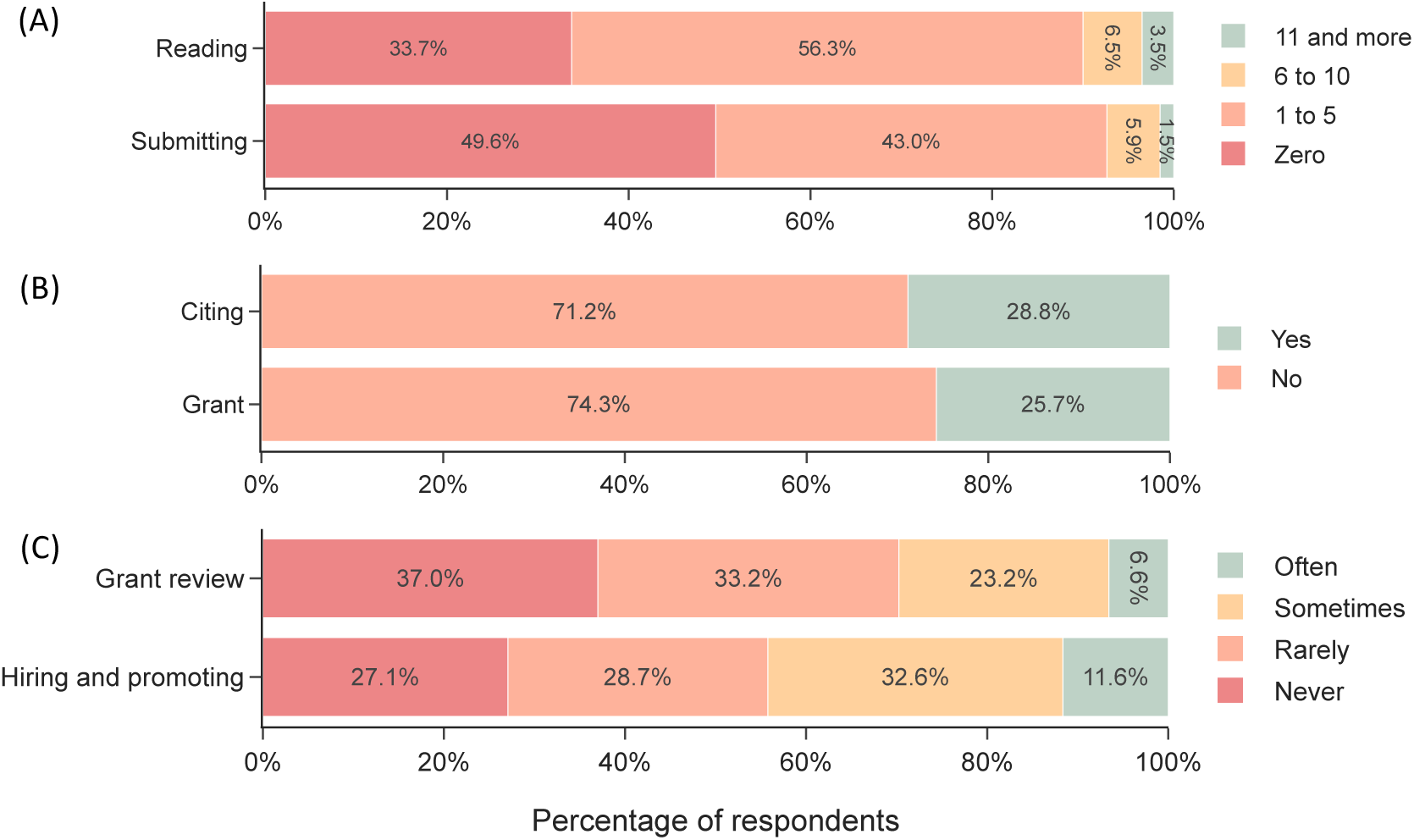
Percentage of respondents by (A) the frequency of reading and submitting preprints per month over the past two years; (B) whether they had cited or included preprints as their representative research achievements in grant applications over the past two years; (C) how frequently they had seen preprints included as the principal investigators’ representative research achievements during grant applications review and the frequency they had seen preprints included as the applicants’ representative research achievements during faculty hiring and tenure promotion. Percentages were calculated based on post-stratification weights.

Our results indicate significant differences across scholar groups. Junior scholars exhibited a higher tendency than senior scholars to read preprints (odds ratio (OR) = 1.463, 95% CI = [1.015, 2.107]), cite preprints (OR = 1.862, 95% CI = [1.298, 2.672]), and include preprints in grant applications (OR = 1.871, 95% CI = [1.246, 2.811]). We also observe substantial disciplinary disparities. Compared with Biomedical and Health Sciences scholars, those in Molecular and Cellular Biology and Quantitative Engineering Biology were more likely to read, cite, and submit preprints and to include preprints in grant applications (see Supplementary **Table 1** for full model results).

### The Pragmatic Engine: Speed over Dialogue and Author Reputation Filtering

Our survey results reveal that reading, citing, and submitting preprints were more like an eagerness for “speed” than a commitment to open science dialogue. As reported by respondents, timeliness was the most influential factor for reading and citing preprints, with 45.3% and 68.3% of respondents selecting it as very or extremely important for reading and citing preprints, respectively (Figure **2A**). Similarly, authors valued preprint utility through speed and ownership. The top motivations for submitting preprints were “gaining immediate visibility and establishing priority” and “accelerating knowledge dissemination,” while “receiving open peer reviews” was the lowest priority, with 75.5% of respondents selecting it as slightly important or not important at all. This suggests that researchers were submitting preprints to broadcast results, not to invite community critique. This is consistent with the low levels of community support among biomedical researchers for the post-publication peer-review platform PubMed Commons, which was shut down after five years of operation from 2013 to 2018 (NLM, 2018). However, the lower priority for open review might stem from the unrealized potential of the preprint system in encouraging dialogue. As one scholar highlighted in a free-text comment: “I’ve done maybe 70 preprints and have received feedback on 2. This has much unrealized potential.” (Man, associate professor, USA, White or Caucasian)

**Figure 2.**
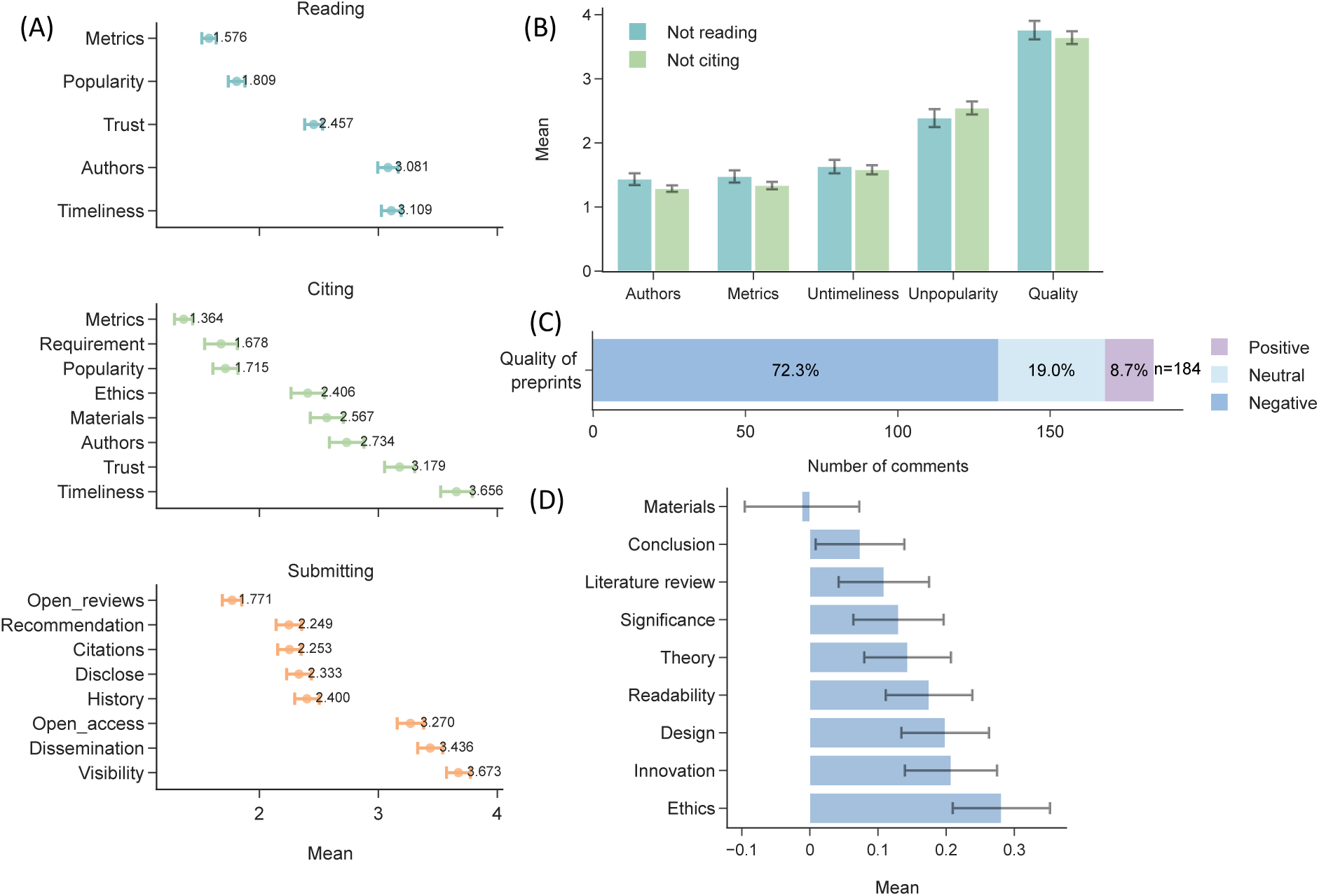
(A) Mean scores of reasons for reading, citing, and submitting preprints reported by respondents. (B) Mean scores of reasons for not reading or citing preprints reported by respondents. (C) Perceived quality (positive, neutral, negative) of preprints in respondents’ free-text comments. (D) Mean scores of the overall quality of preprints perceived by respondents. For panels A and B, means were calculated based on post-stratification weights, with responses coded from 1 (“Not at all”) to 5 (“Extremely”). For panel D, means were coded from −2 (“Poor”) to 2 (“Excellent”). Error bars represent 95% confidence intervals, calculated as the post-stratification-weighted mean ± 1.96 times the weighted standard errors.

Regression results reveal significant differences in scholars’ motivations for engaging with preprints. Compared with senior scholars, junior scholars reported a significantly stronger tendency to cite preprints because of timeliness (OR = 1.735, 95% CI = [1.033, 2.912]) and the popularity of citing preprints in their field (OR = 2.915, 95% CI = [1.724, 4.928]). Respondents in Canada were less likely than those in the United States to cite preprints because of timeliness (OR = 0.569, 95% CI = [0.340, 0.951]). Compared with scholars in Biomedical and Health Sciences, those in Quantitative Engineering Biology showed higher odds of reading preprints because of timeliness (OR = 1.461, 95% CI = [1.077, 1.981]) and citing preprints because of it (OR = 1.568, 95% CI = [1.027, 2.394]). Similar patterns emerged in motivations for submitting preprints. Junior scholars were more likely than senior scholars to submit preprints to increase visibility (OR = 1.514, 95% CI = [1.003, 2.285]). Compared with Biomedical and Health Sciences respondents, those in Molecular and Cellular Biology exhibited higher odds of submitting preprints for visibility (OR = 1.573, 95% CI = [1.084, 2.283]).

Although the need for speed is dominant, users held serious uncertainty about preprint quality, for which reputational heuristics served as a filter to manage risk. Without the traditional “peer review stamp,” uncertainty about preprint quality largely hinged on the acceptance of preprints among potential users. As shown in Figure 2B, among those who reported not reading or citing any preprints over the past two years, 68.6% and 62.4%, respectively, rated concerns about quality as a very or extremely important deterrent. Compared with senior scholars, junior scholars were less likely to report quality concerns as a reason discouraging them from reading (OR = 0.475, 95% CI = [0.283, 0.797]) or citing preprints (OR = 0.576, 95% CI = [0.379, 0.876]). Respondents from Quantitative Engineering Biology were less likely than those from Biomedical and Health Sciences to report preprint quality as a reason discouraging them from reading (OR = 0.505, 95% CI = [0.329, 0.776]). Similar worries emerged from respondents’ free-text comments: among the 184 comments on preprint quality, 72.3% expressed negative views (Figure 2C). The transparency paradox helps explain these concerns: when asked to evaluate different aspects of preprint quality, respondents rated scientific design relatively highly but identified a critical failure in material availability, with 35.4% expressing a negative attitude toward the availability of research materials such as data and code (Figure 2D). To bridge the “need for speed” and “quality uncertainty,” author reputation served as a key filter: among respondents who used preprints, the presence of familiar or notable names in the author list was one of the most influential factors, with 43.9% and 37.1% rating it as very or extremely important for reading and citing preprints, respectively.

These results reveal a dilemma for early-stage researchers. On one hand, they are the most likely to substantially engage with preprints at a vulnerable stage of their career and to value the accelerated pace of scientific communication. On the other, the concerns about preprint quality elicit a heuristic filtering response, filtering by authority, that systematically disadvantages these same researchers. This raises important questions about whether the current evaluation systems mitigate or reinforce this disadvantage with respect to early-career scientists.

### The “Paper Ceiling” and the “Gatekeeper Gap”: Soft Impact vs. Hard Penalties

Our survey indicated that preprints were perceived simultaneously as a networking asset and a career liability in scholars’ professional development. Beyond the use of preprints as readings or references, their influence on scholars’ professional development as academic outputs matters both to individual scholars and to the long-term vision of the preprint ecosystem. As an emerging form of scholarly communication, preprints appear to offer relatively diffuse or indirect career benefits while exposing scholars to more tangible career risks. Respondents viewed preprints as beneficial for fostering research collaborations, enhancing citations, and increasing social impact (Figure **3A**). These benefits were more strongly recognized by junior than senior scholars (collaborations: OR = 2.325, 95% CI = [1.592, 3.396]; citations: OR = 2.045, 95% CI = [1.425, 2.935]; social impact: OR = 2.189, 95% CI = [1.507, 3.181]). Compared with Biomedical and Health Sciences respondents, those in Quantitative Engineering Biology reported more positive effects on collaborations (OR = 1.748, 95% CI = [1.248, 2.450]) and citations (OR = 1.532, 95% CI = [1.112, 2.111]). However, 21.8% of respondents viewed preprints as somewhat or very detrimental to advancement through tenure and promotion, and 38.8% expressed negative views about the impact of preprints on scholars’ career in their free-text answers (Figure **3B**). Echoing the perceived career risk, “lack of recognition in research evaluation” was the leading reason researchers chose not to submit preprints, outweighing fears such as scooping or reputational harm (Figure **3C**). These findings suggest that while many researchers recognize the visibility and accessibility benefits of preprints, concerns about credibility and institutional evaluation pose substantial challenges to their career development. Such implicit penalties function as an important deterrent to preprinting culture.

**Figure 3.**
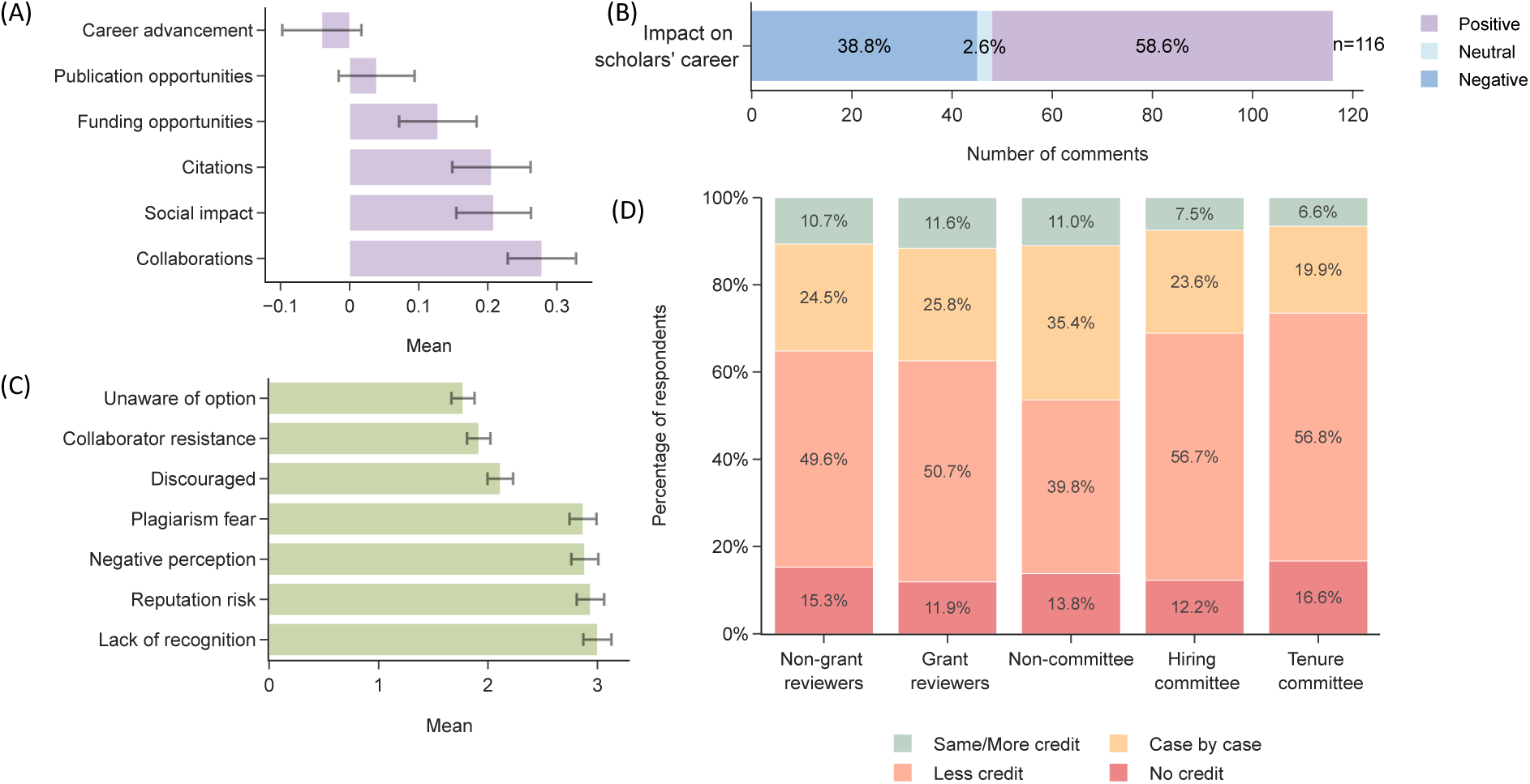
(A) Mean scores of the impact of preprints on their professional development perceived by respondents. (B) Perceived impact (positive, neutral, negative) of preprints on their professional development in respondents’ free-text comments. (C) Mean scores of the reasons for not submitting preprints reported by respondents. (D) Percentage of respondents by how they view the credit preprints should receive in grant applications, tenure job applications and promotion. For panel A, means were calculated based on post-stratification weights, with responses coded from −2 (“Very negative”) to 2 (“Very positive”). For panel C, means were coded from 1 (“Not at all”) to 5 (“Extremely”). Error bars represent 95% confidence intervals, calculated as the post-stratification-weighted mean ± 1.96 times the weighted standard errors. For panel D, percentages were calculated based on post-stratification weights.

Decision-makers’ views across academic evaluation contexts provided a clearer picture of the hard penalties associated with preprints in grant applications, job hiring, and promotion. Among grant reviewers, 29.7% reported sometimes or often encountering preprints as applicants’ research achievements, and 50.7% indicated that they assigned less credit to preprints than to peer-reviewed publications (Figure **3D**). This was consistent with applicants’ experiences, with 49.5% reporting that preprints received less credit during grant applications. Junior scholars were more likely than senior scholars to support giving preprints more credit in grant applications (OR = 2.149, 95% CI = [1.363, 3.389]) and to report giving them higher credit as grant reviewers (OR = 2.503, 95% CI = [1.273, 4.924]). The pattern was similar in tenure, hiring, and promotion: 44.2% of committee members sometimes or often observed preprints listed as candidates’ research achievements, but 56.7% assigned less credit to preprints in faculty job applications and 56.8% did so in tenure reviews. Only 7.5% and 6.6%, respectively, gave preprints the same or more credit.

The overall conservative attitude reflects the consideration of risk management and efficiency among academic gatekeepers. As stated in respondents’ free-text comments: “Nobody has time to read preprints from 30 candidates for a position or award to determine their value. Thus, we use journal publications. At least as a reviewer, we know there has been some bar surpassed.” (Man, associate professor, USA, White or Caucasian) However, compared with committee members, respondents who had not served on hiring or promotion committees displayed a clear divergence, with substantially greater openness to case-by-case evaluation (35.4%) rather than applying a blanket penalty. This suggests a structural challenge to preprint adoption: gatekeepers are systematically more likely to discount this form of contribution to the scientific literature. This attitude may be self-reinforcing if it selects out those scientists prioritizing rapid communication through preprints.

### Perceived impact on science and society: Accelerating openness and sharing versus fraudulence and emerging AI threats

Respondents held overall conservative perceptions of the impact of preprints on the scientific community and society. When discussing potential positive impacts, respondents showed stronger agreement with promoting open access to research for all audiences and accelerating publishing and knowledge sharing. Overall, 49.1% and 40.2% of respondents, respectively, agreed very or extremely strongly with these contributions (Figure **4A**). In contrast, reducing reliance on traditional top-tier journals and driving changes in research evaluation criteria received less emphasis. When asked about potential negative impacts, respondents expressed greater concern about spreading misleading or fraudulent information and harming research integrity through papers created using large language models, with 44.0% and 38.9%, respectively, agreeing very or extremely strongly (Figure **4B**). These concerns outweighed wasting readers’ time on unvetted studies and weakening public trust in science. Weighted regression results indicate significant differences by academic rank, country, and field. Especially, junior scholars reported more positive perceived impacts of preprints and less concern about their negative effects (see Supplementary **Note 3** for full results).

**Figure 4.**
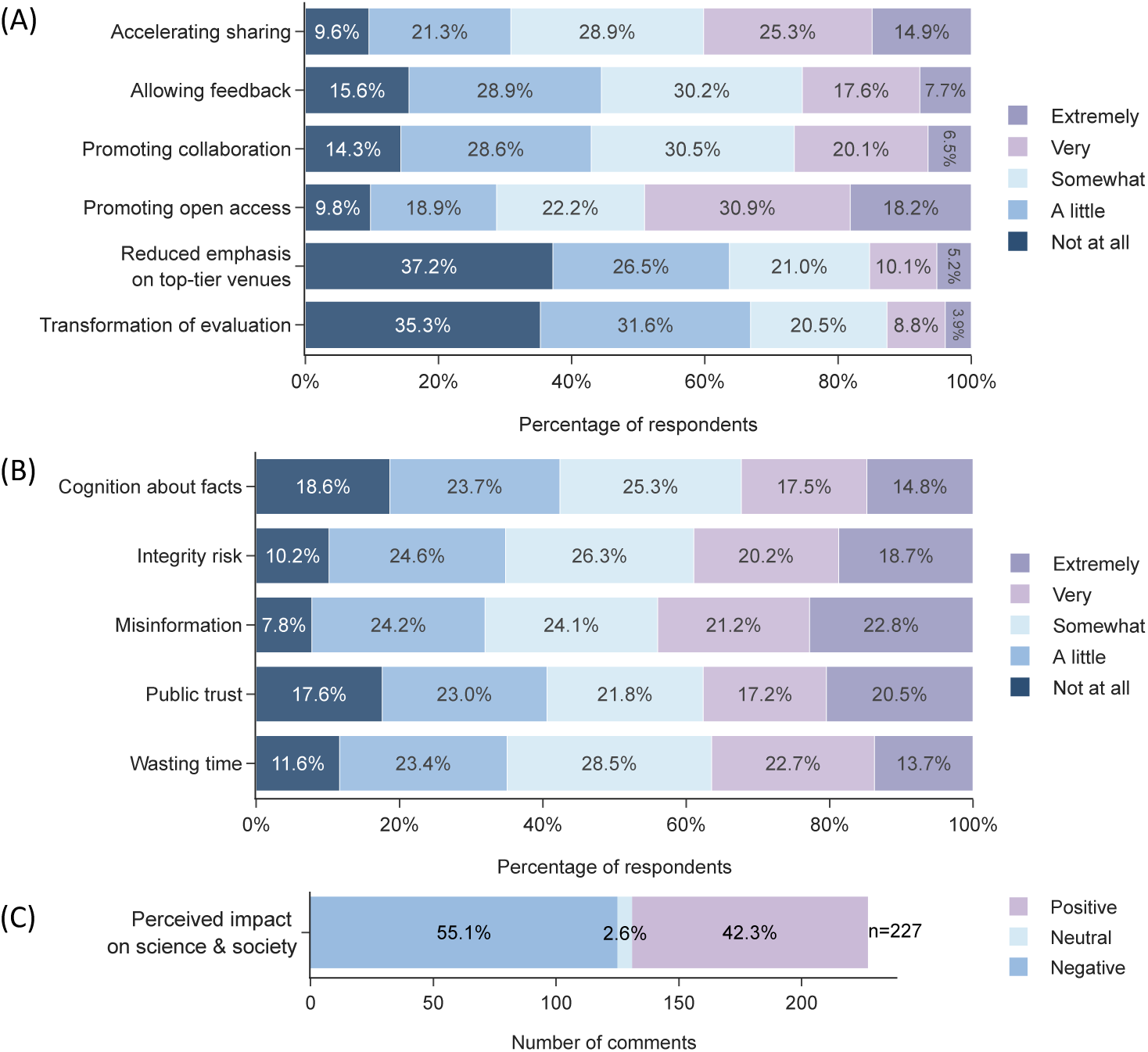
Percentage of respondents by: (A) how they view each statement about the positive impact of preprints. (B) how they view each statement about the negative impact of preprints. (C) their sentiment toward the perceived impact of preprints on the scientific community and society in free-text comments. Percentages were calculated based on post-stratification weights.

Sentiment labeling of the free-text comments offers deeper insight into how scholars perceive the positive and negative impact of preprints on the scientific community and broader society. Among the 227 free-text answers, 55.1% expressed negative views about the impact of preprints on science and society, 42.3% expressed positive views, and 2.6% were neutral (Figure **4C**). Positive views most often emphasized that preprints help alleviate pressures associated with the relatively slow peer-review process, enable faster and cost-free dissemination through open access, and facilitate more timely and transparent feedback, dialogue, and potential collaboration. As respondents noted:

> “*Open access of preprints lets me comfortably not pay for Gold Open Access fees for journal publications. I use that money to pay undergraduate researchers instead*.” (Woman, assistant professor, USA, White or Caucasian)
>
> “*They are so helpful in reducing the HUGE burden that the publishing industry takes on my workload…Hopefully this will encourage journals to be faster and to require fewer additional experiments just because a reviewer asks for them.*” (Other gender, associate professor, USA, other race)
>
> “*But I believe that if a field does use them, the fact that one can get feedback from multiple peers not just the chosen peer reviewers is a positive. If I was to evaluate someone’s record and they listed pre-prints I would go look at the feedback*.” (Man, full professor, USA, White or Caucasian)
>
> “*I primarily use preprints as help sharing my work within my scientific network. I find they’re most useful as a tool for communicating a complete scientific story to individual colleagues and potential collaborators*.” (Woman, assistant professor, USA, White or Caucasian)

On the other hand, negative sentiments came from the concerns on fraudulence and misinformation in the context of low public trust in science and AI threats to scientific integrity and quality, the damage to the (blind) peer-review system, and the potential exacerbation of the Matthew Effect caused by preprints:

> “*It is entirely possible to have fully fabricated fictitious documents uploaded. This can affect real life… In medicine or any other field with great importance for the general population they are not useful, and they can be a source of misinformation that may risk for the population safety*.” (Man, full professor, Canada, Hispanic or Latino)
>
> “*My concerns about them have been magnified by AI. I worry we will see professors mass generating pre-prints with AI and listing them and then generating more pre-prints that cite the previous pre-prints…Not only would there be AI generated garbage that students or the public could find and mistakenly believe to be legitimate, but it could allow some people to game sites like Google Scholar or ResearchGate and crowd out legitimate scholars who are publishing at a slower pace because they are actually doing real studies and they are going through peer review.*” (Man, associate professor, USA, White or Caucasian)
>
> “*Preprints undermine the blind peer review process (by revealing authors identities) and create a potentially dangerous situation where members of the public do not realize that the preprint is not peer reviewed. I think they should be banned*.” (Woman, full professor, USA, Hispanic or Latino)
>
> “*They exacerbate halo effects and this would be worsened if preprints became more common. Preprints would exacerbate inequalities between low profile labs in developing nations and US high profile institutions*.” (Man, full professor, USA, other race)

These results point to an unmet need in the preprint ecosystem: accountability mechanisms. At present, integrity risks, misinformation, and resultant trust are handled in the publishing ecosystem through editorial and review mechanisms that have no counterpart in preprints.

## Discussion and conclusions

Drawing on a large-scale survey of US and Canadian biomedical scholars, this study reveals a broadening landscape of preprint adoption aimed at accelerating scientific communication, alongside significant structural barriers arising primarily from concerns about preprint quality and perceived penalties in evaluation.

Our study reveals that the use of preprints exhibits an emphasis on “speed” rather than “dialogue,” which limits the role of preprints in accelerating open science. As a rapidly growing channel for the timely and public dissemination of scientific findings, preprints represent one of the most significant developments in the broader movement toward open science, addressing the limitations of traditional publishing in open access and fast visibility. In this context, prioritizing speed is understandable given the intensifying competition surrounding publication and authorship. Nevertheless, the strong pragmatic focus on speed may constrain the broader potential of preprints to advance open science through sharing and dialogue. Therefore, the preprint ecosystem needs to move beyond speed alone and further promote openness and knowledge sharing by cultivating a widely recognized culture of academic communication and dialogue.

Although preprints are increasingly used as evidence of research achievement by scholars, the penalties on preprints in academic evaluation may undermine both scholars’ career and preprinting culture, therefore necessitate reforming the future academic evaluation system. The penalties on preprints are driven largely by concerns about risk and efficiency from gatekeepers’ perspective. As gatekeepers of science, reviewers and committee members must effectively manage risk when allocating funding and positions, for which the traditional peer-review system has historically served as a reliable and efficient quality-verification mechanism. However, it is foreseeable that preprints will become an increasingly established component of scholarly achievement, and the resistance to recognizing them may exacerbate the conflict and misalignment in evaluation. If the evaluation system is to adapt, a more comprehensive evaluation guideline should be developed to better integrate preprints into it.

Concerns about preprint quality stem in part from the unavailability of research materials, such as data and code. Although such unavailability may arise for reasons such as data disclosure restrictions or authors’ concerns about criticism, readers nonetheless perceive it as a signal of uncertainty and may instead seek the authors’ names and reputation for identifying reliable information. This contrasts with evidence from earlier years, which suggested that author information played a relatively limited role in shaping trust in preprints (Soderberg et al., 2020). As the preprint ecosystem expands, this reliance on reputation-based filtering may reinforce the ‘Matthew Effect’: well-known scholars gain disproportionate visibility, while junior researchers become more easily overlooked. Additionally, author or institutional reputation may not be a reliable indicator of preprint quality, as even established researchers and prestigious institutions can produce work of variable quality.

More concerningly, both structured and free-text answers highlight the worries about preprints on spreading misleading or fraudulent information, especially in the context of the growing use of AI-assisted writing and research. The rise of AI-assisted content creation has intensified concerns about preprints, as it enables the production of a flood of papers containing little new knowledge or even fraudulent information (Boboris, 2025; Watson, 2025). With the rapid growth and fast dissemination of preprints, some premature or low-quality research may widely circulate unwarranted scientific claims via social media, which, if later shown to be incorrect, would further reduce public trust in science (Guterman & Braunstein, 2020; Janda et al., 2022; J. Teixeira da Silva, 2017; Ravinetto et al., 2021).

These tensions highlight structural issues with the adoption of preprints in biomedicine that arise from a common cause: quality concerns. These findings highlight the lack of better solutions to signaling rigor than author reputation. This points to an unmet need: a scalable quality summarization framework able to communicate key quality features of preprint submissions, especially on methodological transparency and rigor, through standardized indicators such as badges for data and code availability, signals of transparent research practices, links to peer-reviewed versions, and visible community reviews. This would serve as a vital safeguard and a signaling mechanism for both authors and readers. Furthermore, our results suggest that mechanisms to signal methodological rigor and materials availability would disinhibit other aspects of the preprint movement like personnel gatekeeping and junior scientist disadvantage. By mitigating structural barriers identified here, such as filtering by authority and personnel selection, preprints’ role as a scientific accelerator would be much more effectively realized.

This study has limitations. Although we have adopted post-stratification weights by country and field, they may not adjust for the potential nonresponse bias associated with other factors such as academic rank, institution type, demographic characteristics, or prior preprint use. The findings on the quality and impact of preprints mainly reflect respondents’ subjective perceptions, which highlights the need for future examination based on more rigorous empirical evidence. The results should therefore be interpreted as reflecting the views of the surveyed biomedical scholars in the United States and Canada.

## Supplementary file

### Supplementary Note 1: Post-stratification Weighting

Post-stratification weights are constructed from eight strata defined by crossing four broad disciplinary fields with two countries. For a given discipline-by-country stratum h, Nₕ and nₕ denote the population and survey-sample counts, respectively, while N and n denote the total population and sample sizes. The raw weight is the ratio of the population share to the survey sample share:

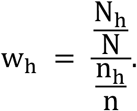

Weights are then normalized to have a mean of 1:

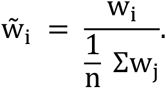

The normalized respondent-level weights are applied to descriptive analysis and to the ordinal logistic regression models, thereby aligning the analytic sample with the population’s joint discipline-by-country distribution.

### Supplementary Note 2: Statistical modeling

We use ordinal logistic regression to examine group differences in attitudes toward, and perceptions of, preprints. The outcome variables include the frequency of reading, citing, and submitting preprints; respondents’ reasons for (not) engaging in these practices; perceived impacts of preprints on the scientific community, society, and scholars’ professional development; and the extent to which preprints are perceived or granted credit as research achievements. Academic rank, gender, country, and discipline are included as predictors to assess potential differences in tendencies across groups. The equations of ordinal logistic regression are as follows:

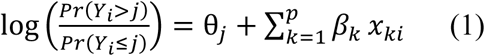

In Equation 1, *Yi* denotes the ordinal level of an outcome for respondent *i*, whereas *j* represents a threshold between the *n* categories of the outcome variable (1 < *j* < *n* − 1). *Pr*(*Y_i_* ≤ *j*) and *Pr*(*Y_i_* > *j*) refer to the probability of being at or below *j* and being above *j*, respectively. 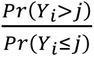 denotes the odds of being above versus at/below *j*. θ_j_ is the baseline cumulative log-odds at *j*. *x_k_* is the *k*th’ predictor for respondent *i*. *β_k_* is the coefficient of *x_k_* in the model. Taking the logarithm of the both sides of Equation 1, 1-unit increase of *x_k_* will multiply the odds 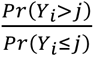 by *ext*(*β_k_*), as shown in Equation 2:

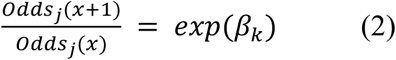

In the manuscript, we report the odds ratio *exp*(*β_k_*) rather than original *β_k_*.

**Supplementary Table S1:**
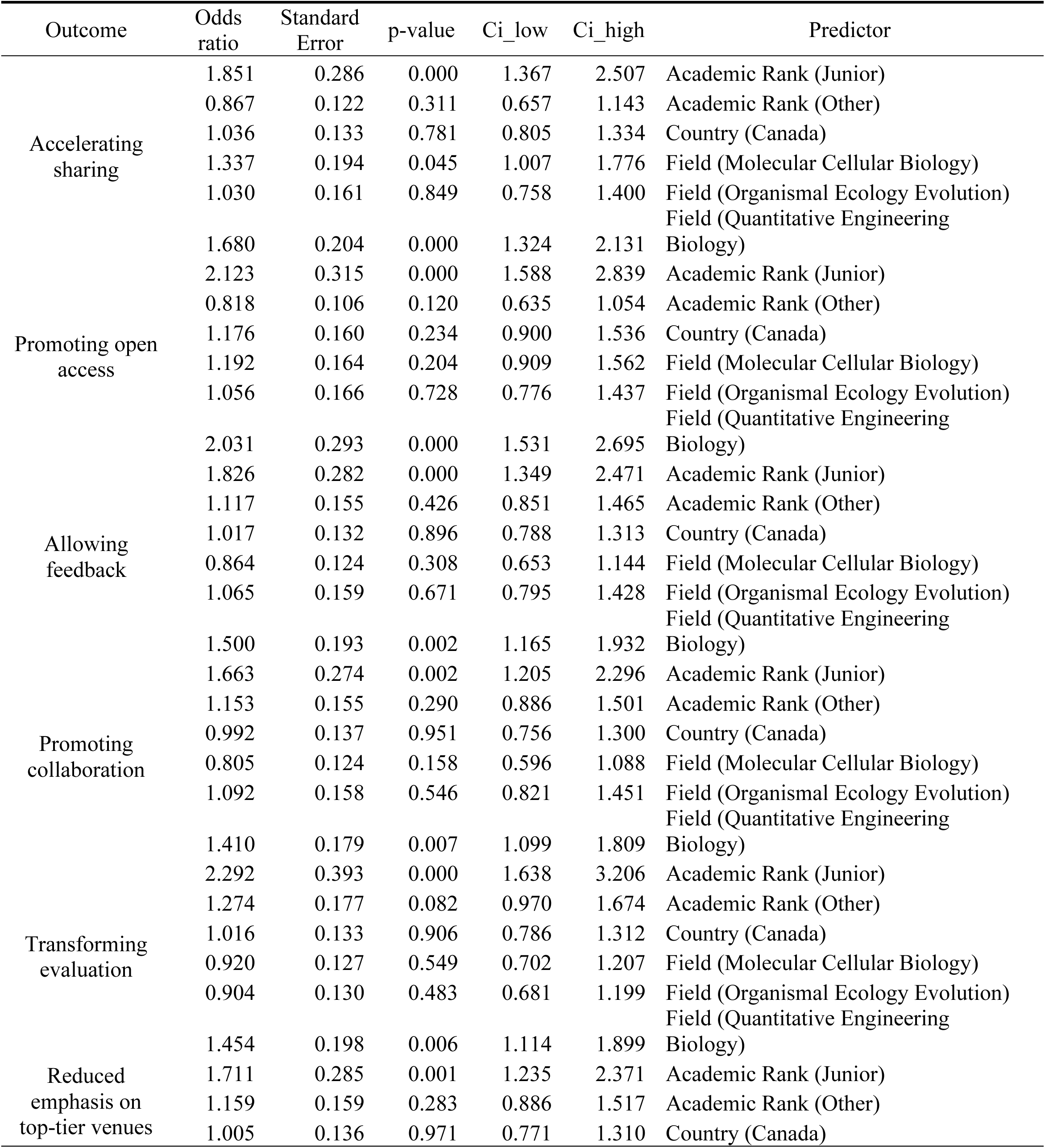
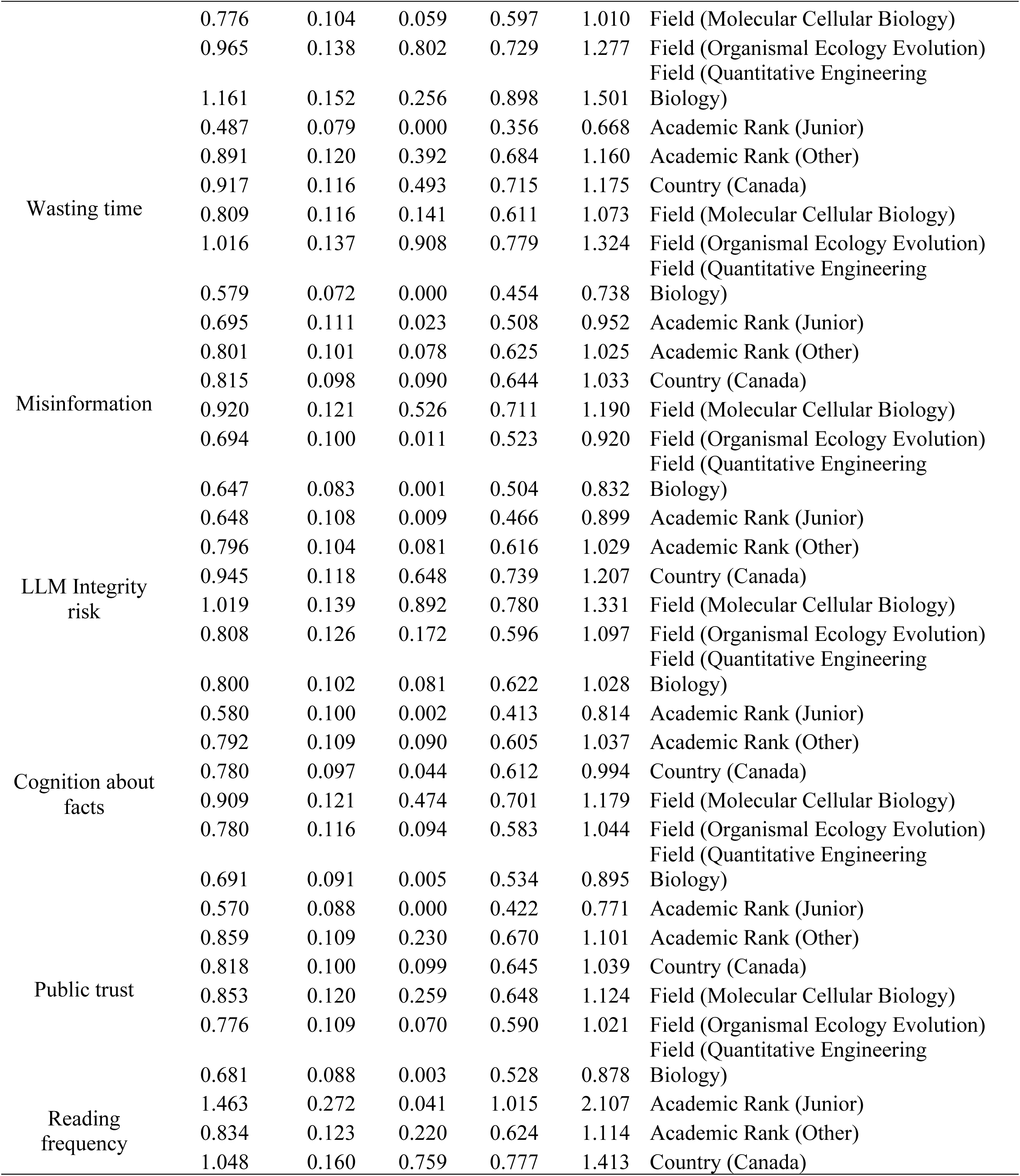
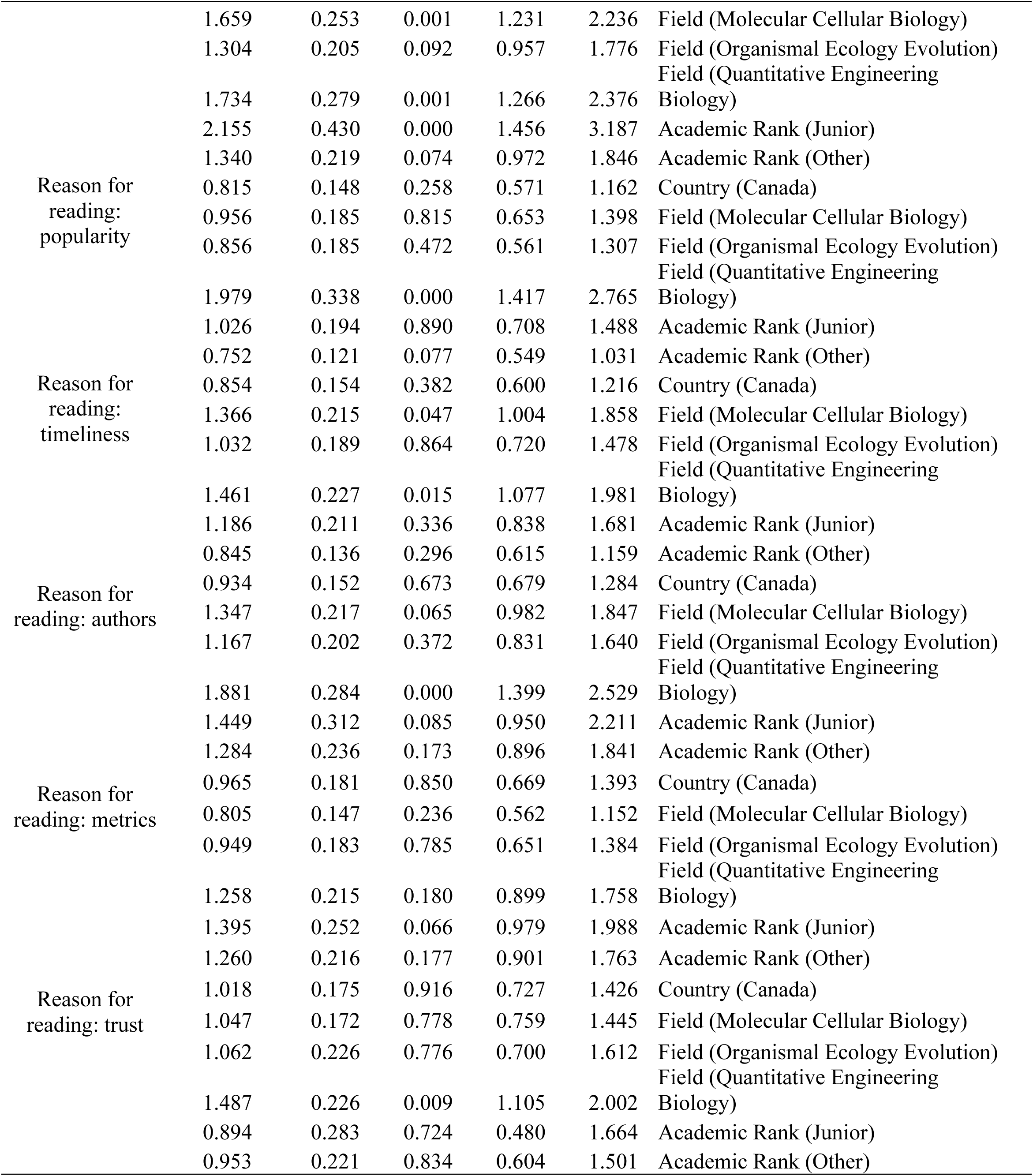
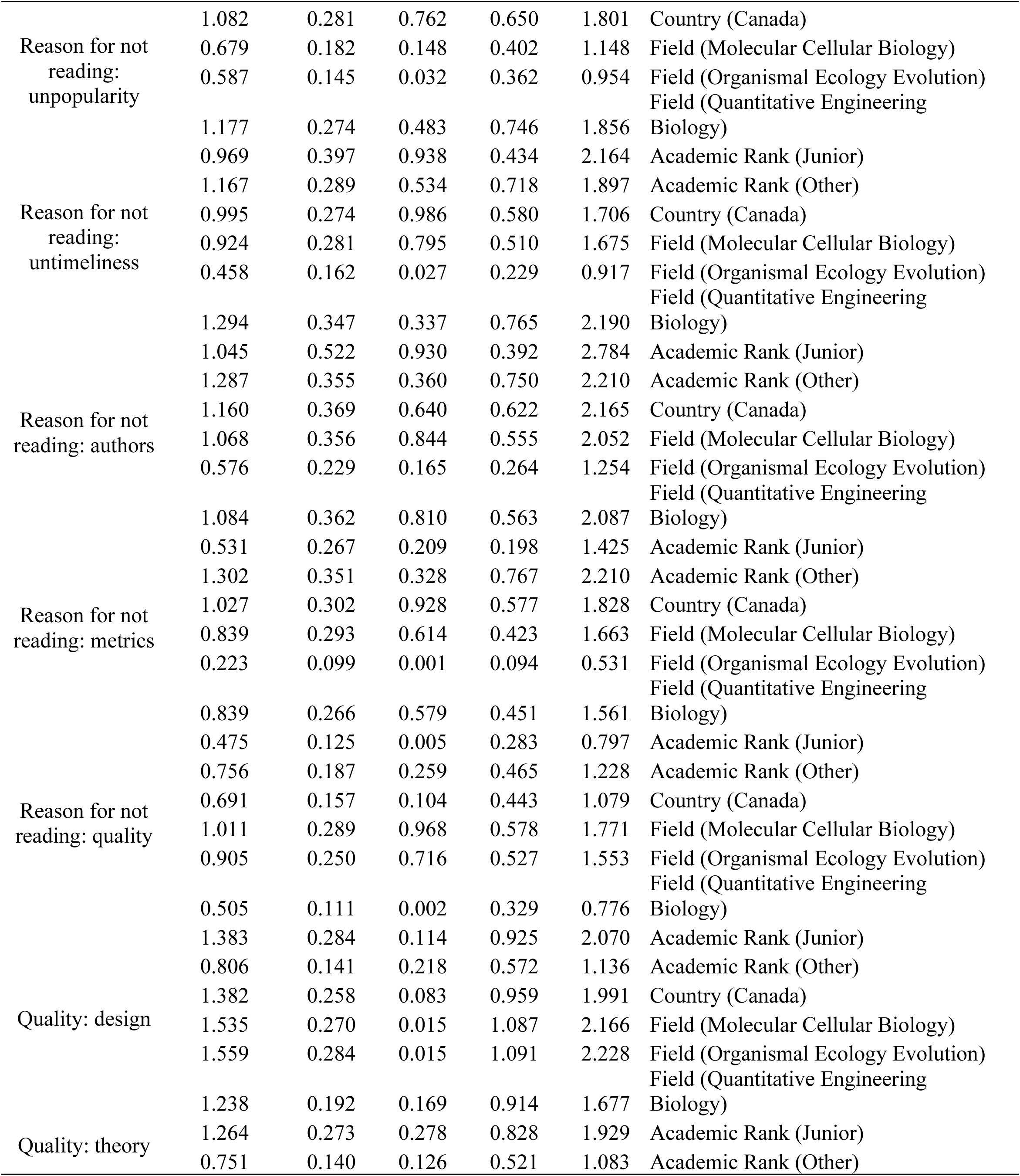
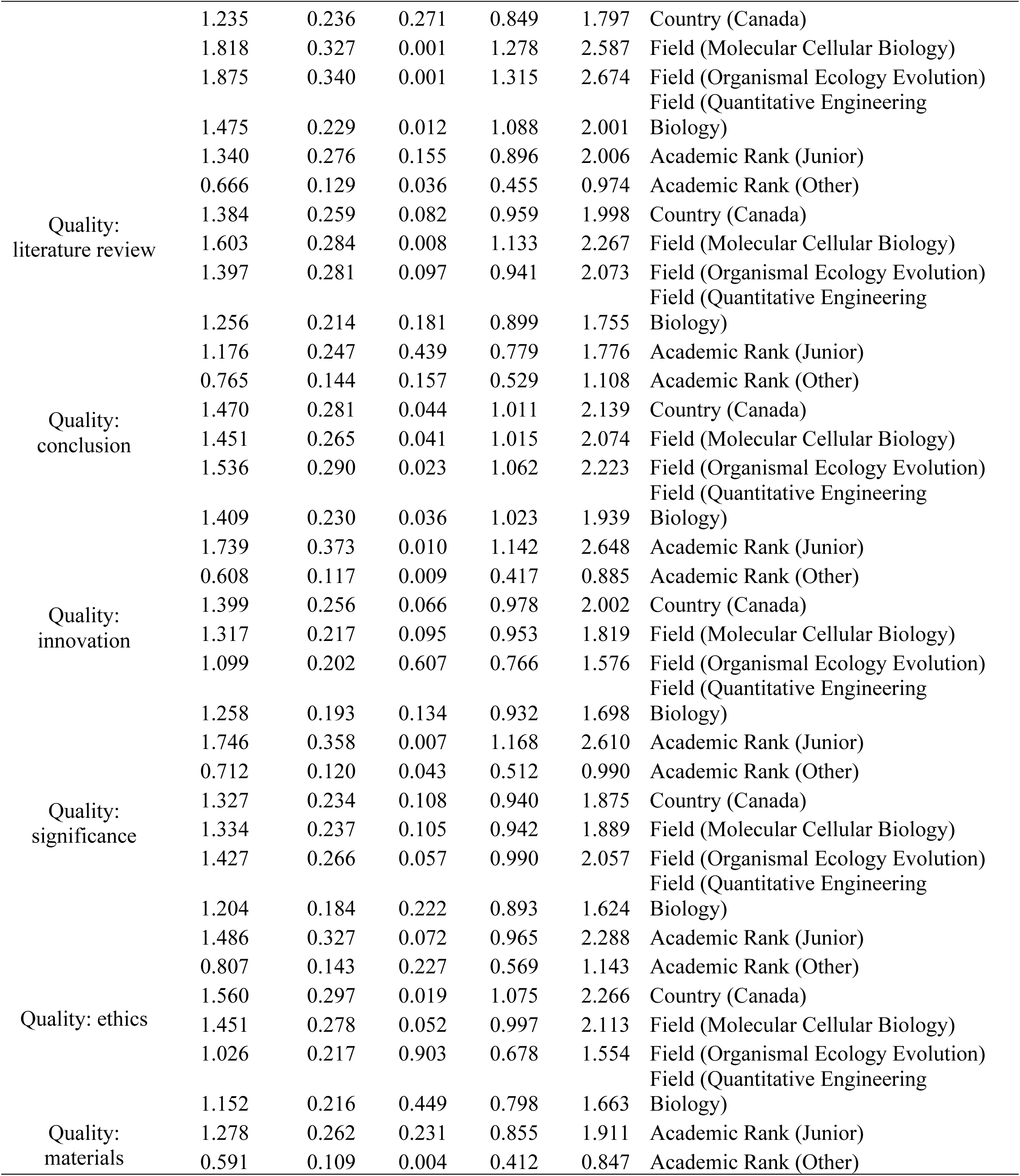
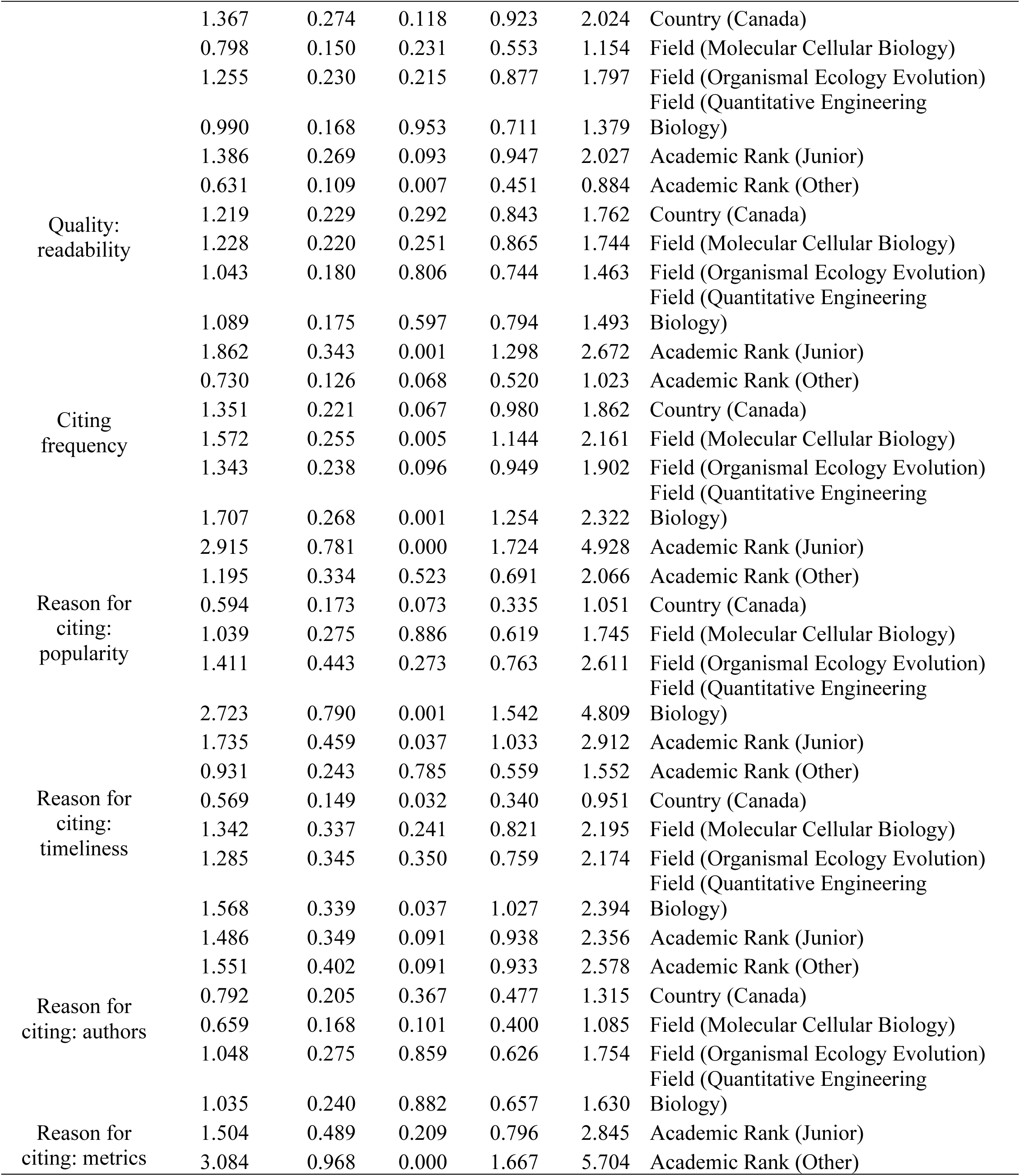
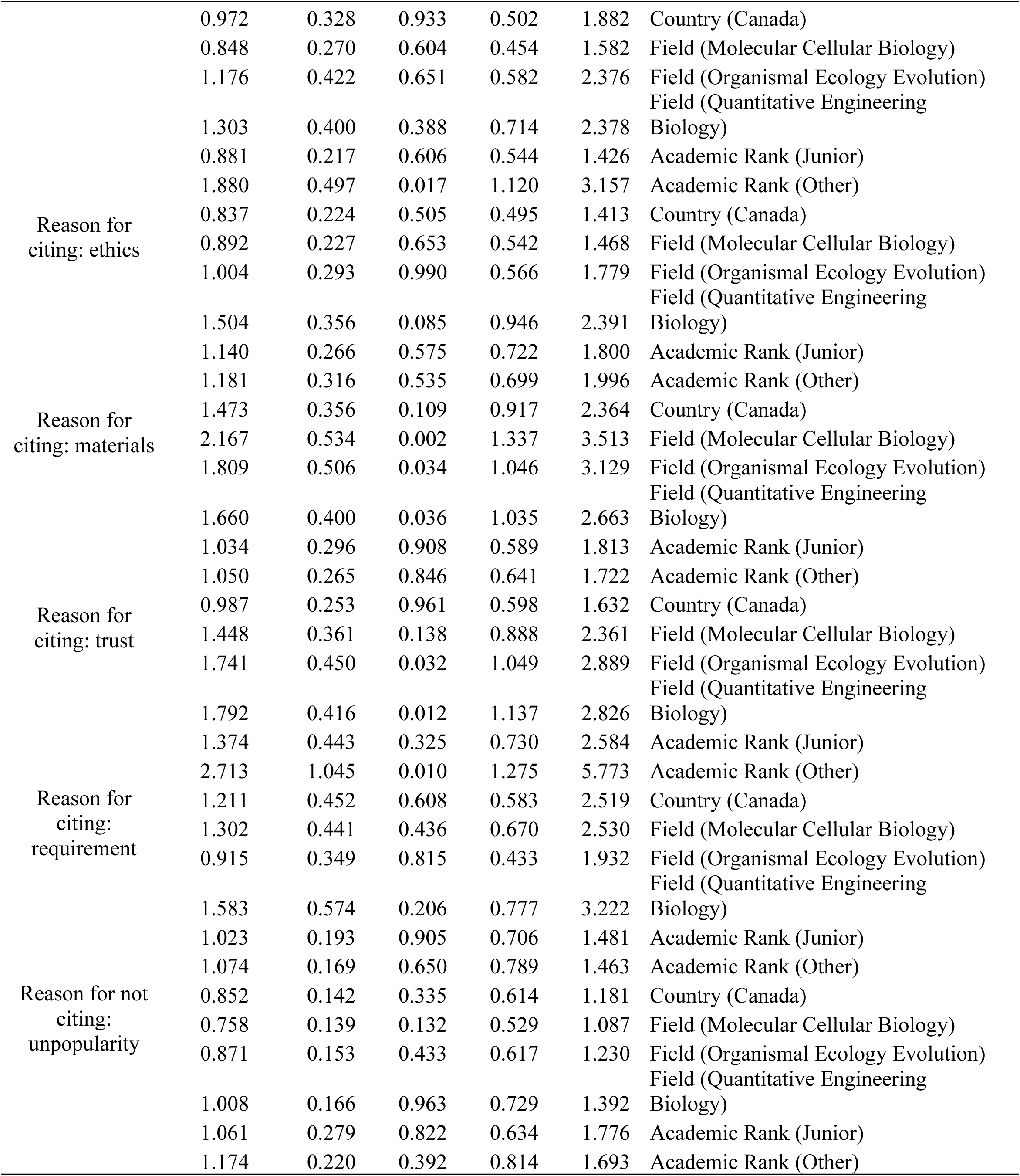
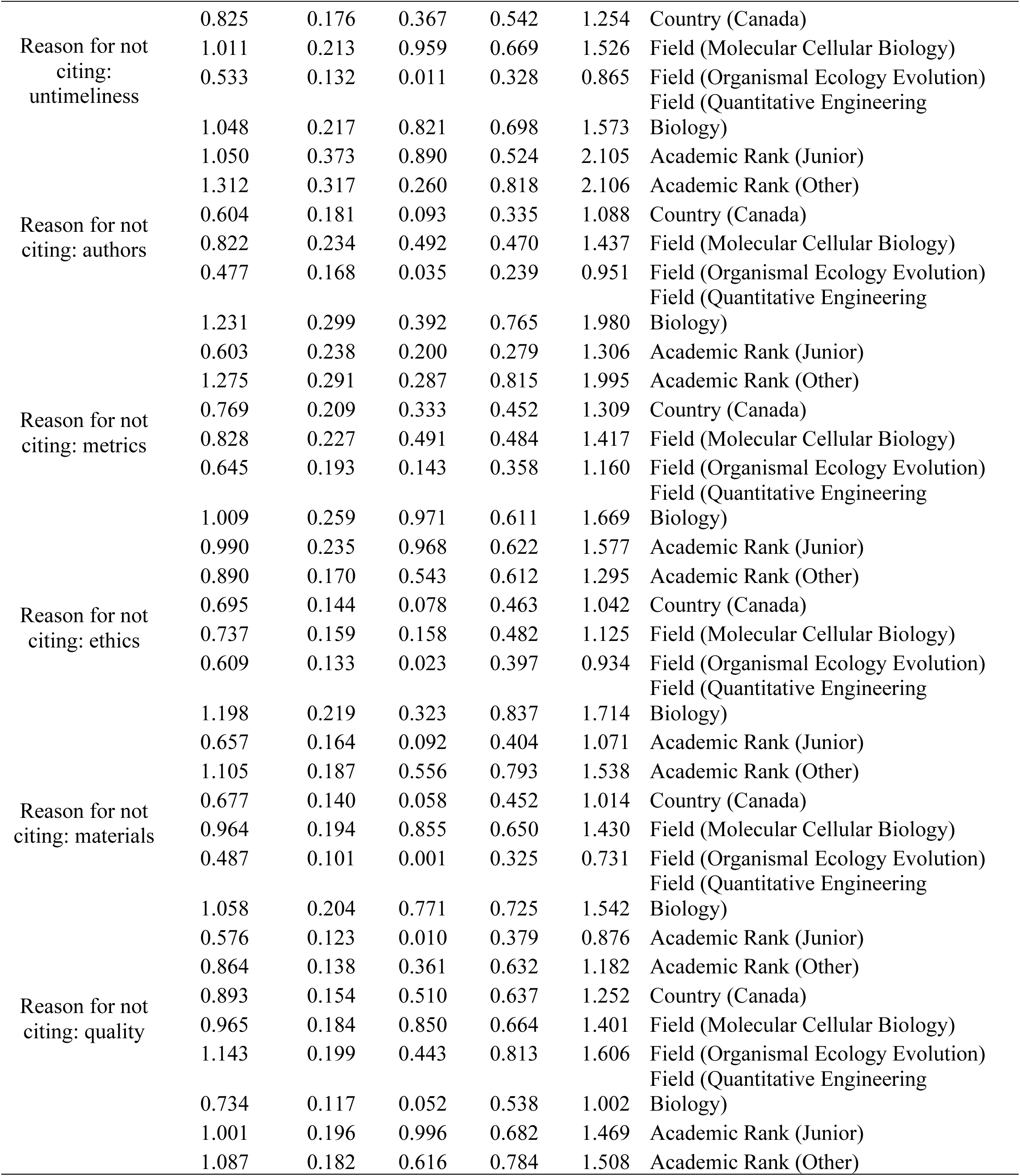
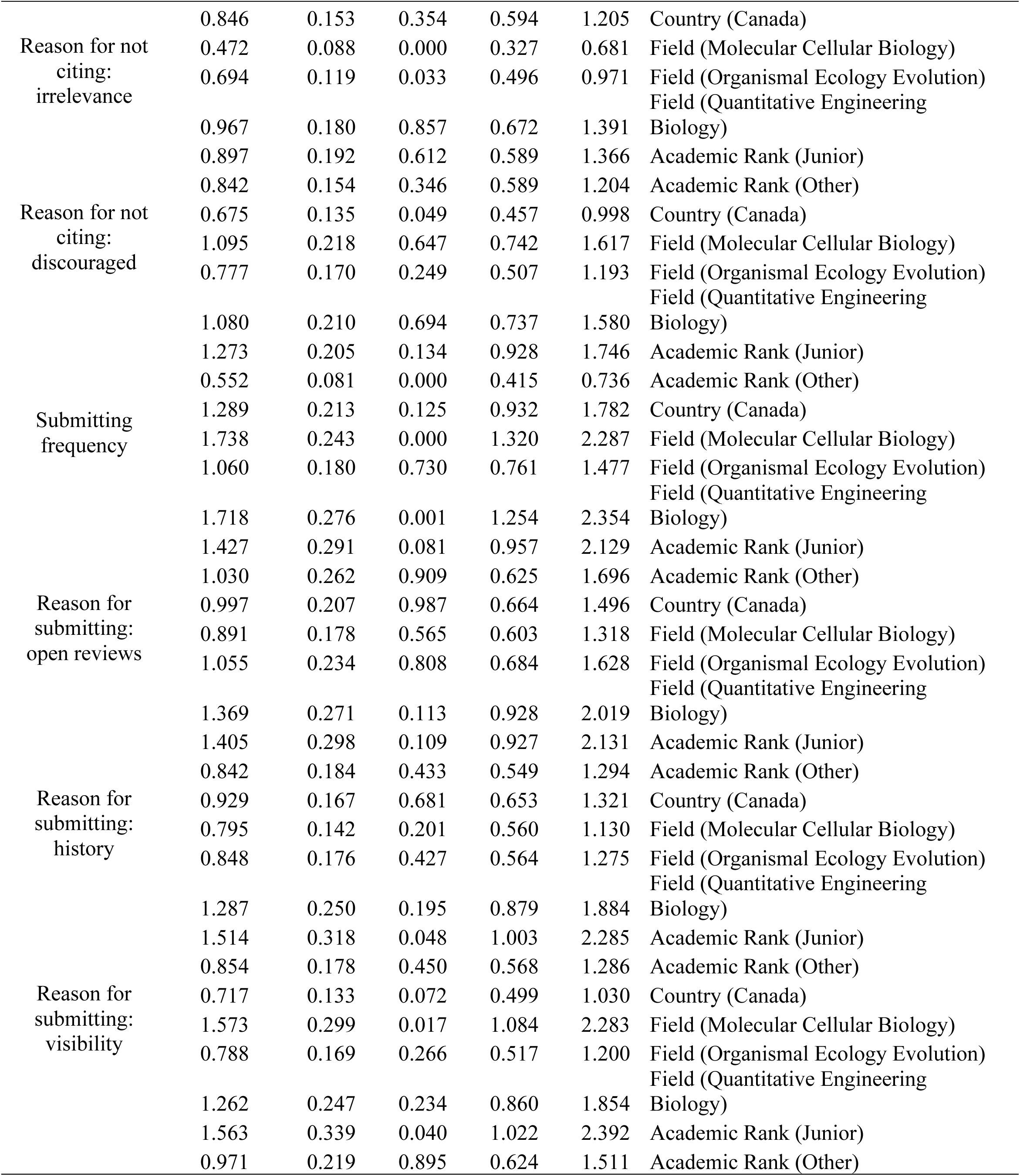
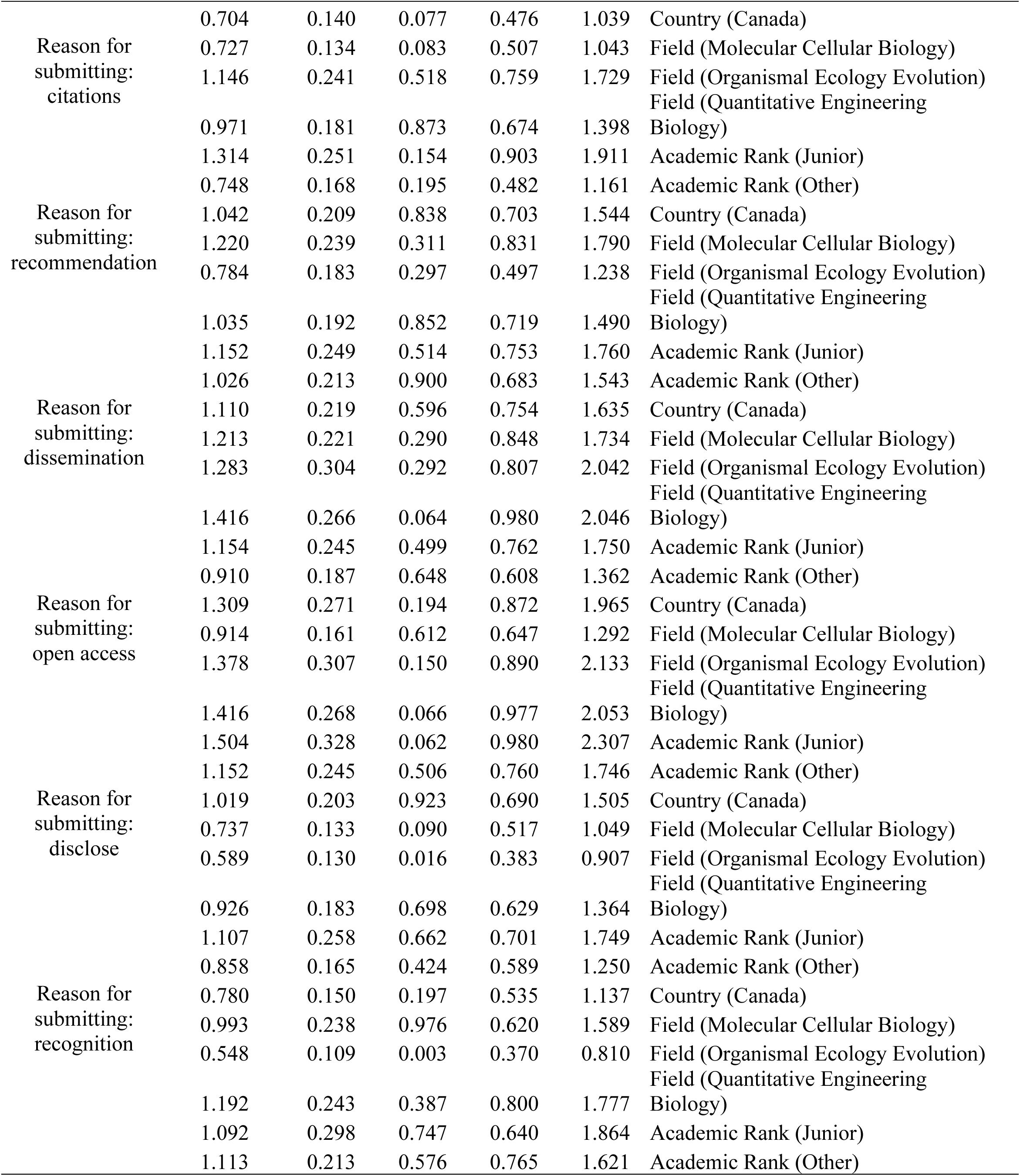
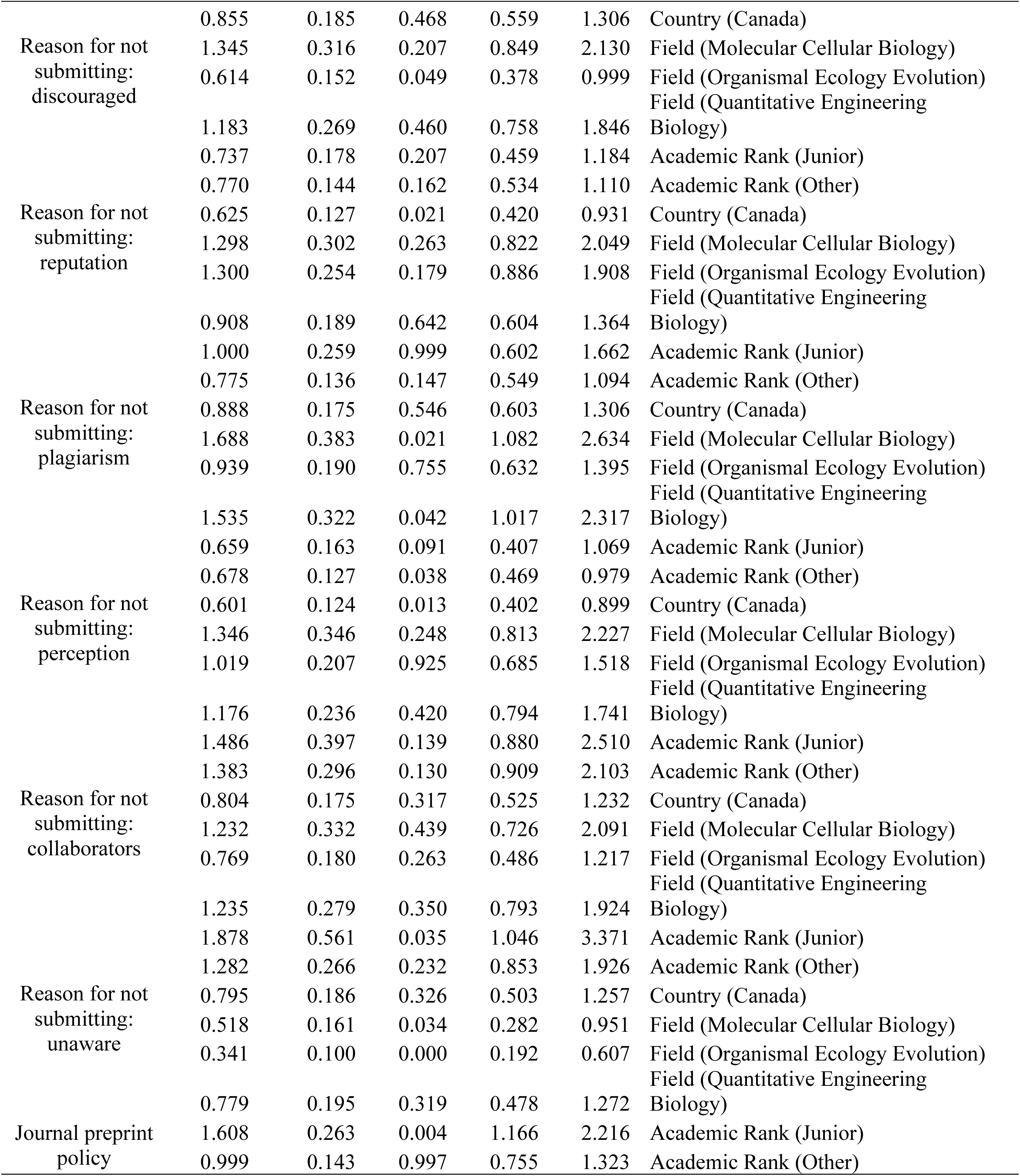
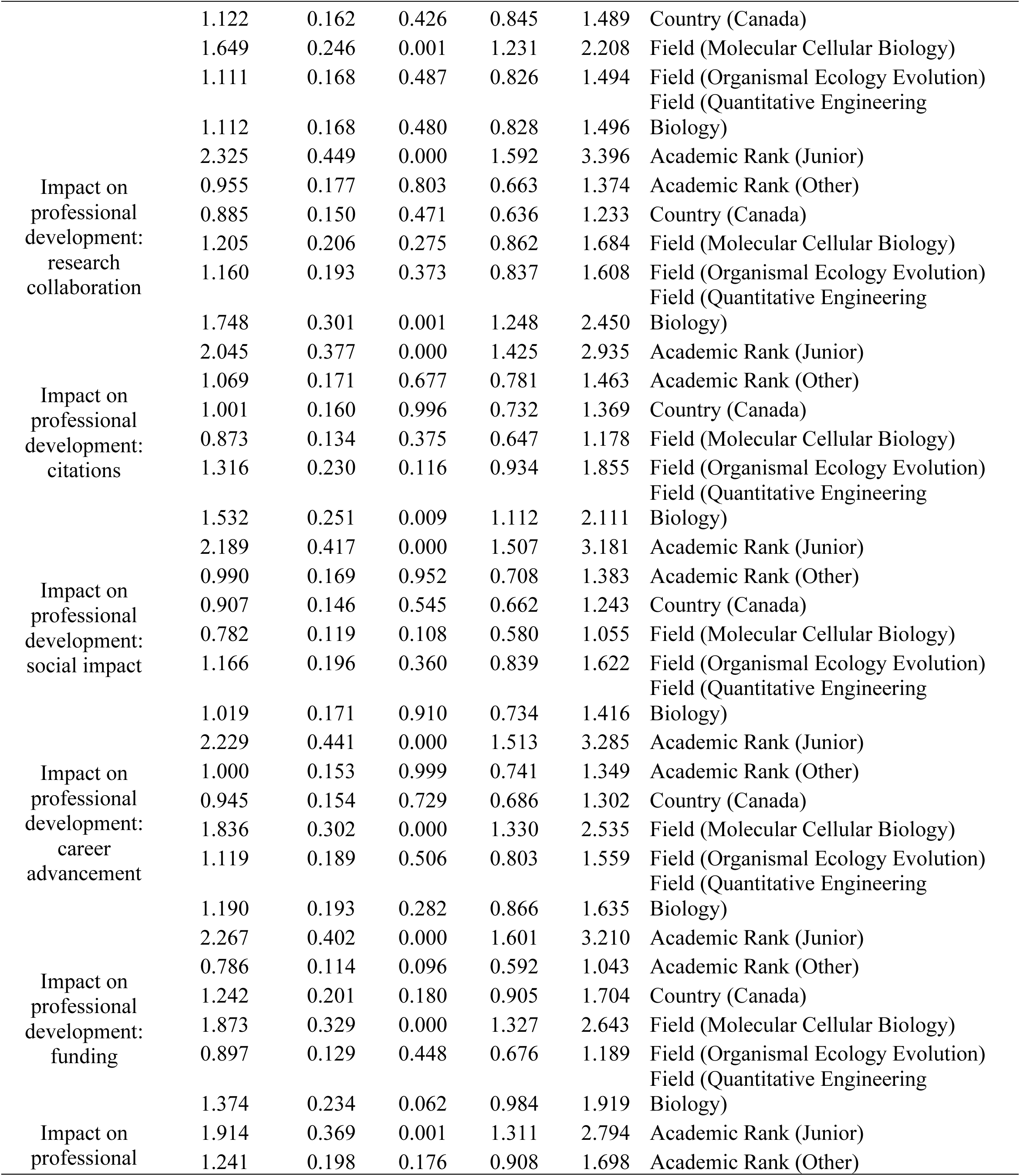
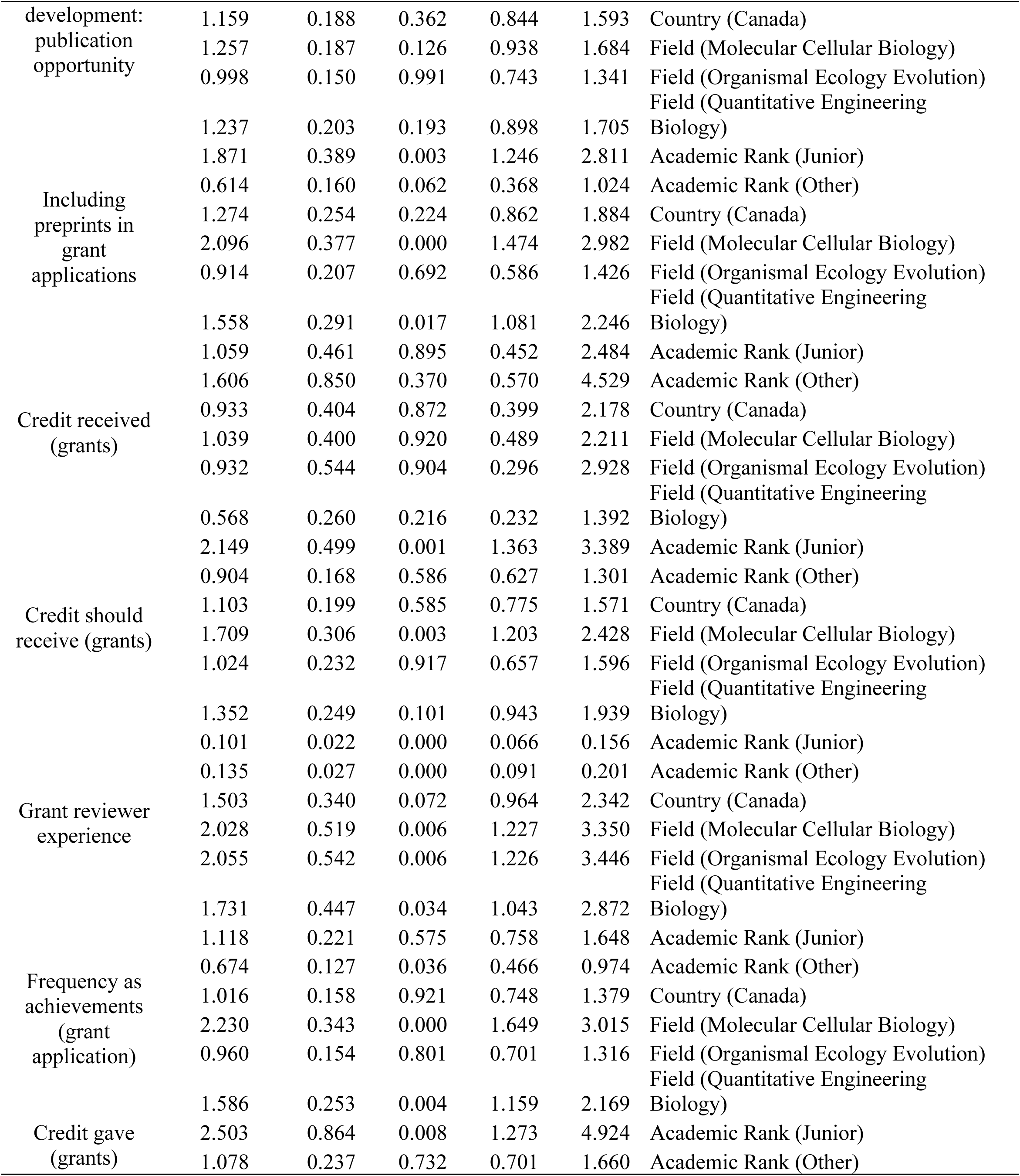
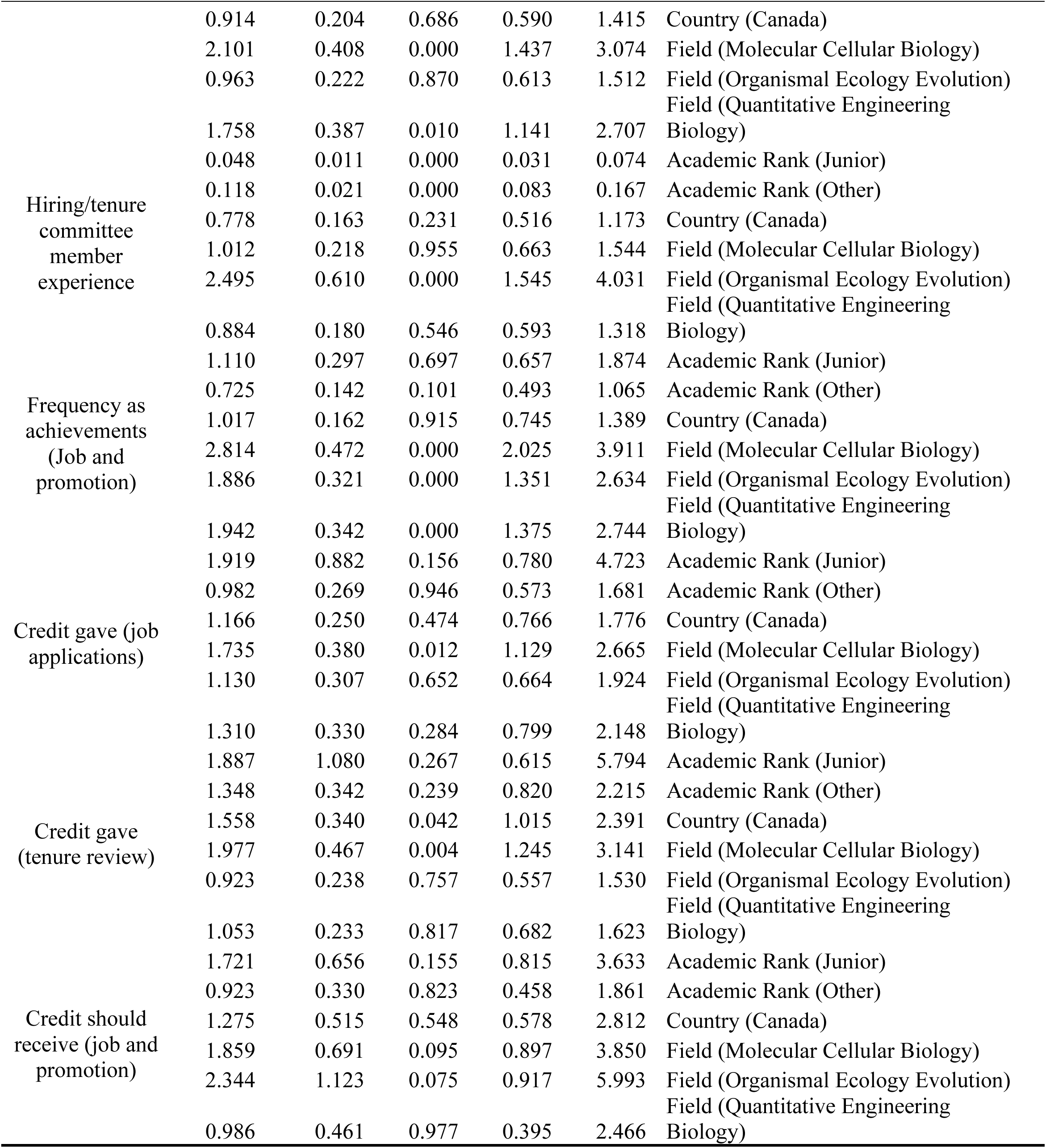
Ordinal logistic regression results.

### Supplementary Note 3: Full ordinal regression results of Section [Perceived impact on science and society]

Compared with senior scholars, junior scholars report substantially more positive perceived impacts of preprints on the scientific community and society, and less concern about negative influences. Specifically, junior scholars exhibit higher odds of agreeing that preprints accelerate publishing and knowledge sharing (OR = 1.851, 95% CI [1.367, 2.507]), promote open access (OR = 2.123, 95% CI [1.588, 2.839]), allow for broader community feedback on research (OR = 1.826, 95% CI [1.349, 2.471]), strengthen collaboration (OR = 1.663, 95% CI [1.205, 2.296]), drive the transformation of research evaluation criteria (OR = 2.292, 95% CI [1.638, 3.206]), and reduce the emphasis on traditional publishing venues like top-tier journals (OR = 1.711, 95% CI [1.235, 2.371]). In contrast, junior scholars expressed lower odds of endorsing negative views, including wasting readers’ time on unvetted studies (OR = 0.487, 95% CI [0.356, 0.668]), malicious use (OR = 0.695, 95% CI [0.508, 0.952]), harming research integrity by publishing papers created using tools like large language models (OR = 0.648, 95% CI [0.466, 0.899]), harming people’s cognition about things or facts (OR = 0.580, 95% CI [0.413, 0.814]), and weakening public trust in science (OR = 0.570, 95% CI [0.422, 0.771]). Disciplinary differences are identified with Biomedical and Health Sciences as the baseline. Respondents in Molecular and Cellular Biology were more likely to perceive preprints as accelerating publishing and knowledge sharing (OR = 1.337, 95% CI [1.007, 1.776]). Perceptions were especially favorable in Quantitative Engineering Biology, where respondents reported higher odds of viewing preprints as accelerating publishing and knowledge sharing (OR = 1.680, 95% CI [1.324, 2.131]), promoting open access (OR = 2.031, 95% CI [1.531, 2.695]), enabling feedback (OR = 1.500, 95% CI [1.165, 1.932]), strengthening collaboration (OR = 1.410, 95% CI [1.099, 1.809]), and driving transformations in evaluation (OR = 1.454, 95% CI [1.114, 1.899]). At the same time, scholars in this field expressed fewer worries about the negative impact of preprints: Quantitative Engineering Biology respondents were less likely to view preprints as a waste of readers’ time on unvetted studies (OR = 0.579, 95% CI [0.454, 0.738]), less likely to worry about malicious use (OR = 0.647, 95% CI [0.504, 0.832]), harming people’s cognition (OR = 0.691, 95% CI [0.534, 0.895]), or reduced public trust (OR = 0.681, 95% CI [0.528, 0.878]). A similar pattern appeared for Organismal and Ecology Evolution, where respondents had lower odds of concern about malicious use (OR = 0.694, 95% CI [0.523, 0.920]). Canadian respondents also had lower odds of concern about harming people’s cognition (OR = 0.780, 95% CI [0.612, 0.994]).

